# The Evolutionary History and Impact of Bacterial tRNA Modifications

**DOI:** 10.1101/251322

**Authors:** Gaurav D Diwan, Deepa Agashe

## Abstract

Along with tRNAs, enzymes that modify anticodon bases are a key aspect of translation across the tree of life. tRNA modifications extend wobble pairing, allowing specific (“target”) tRNAs to recognize multiple codons and cover for other (“non-target”) tRNAs, often improving translation efficiency and accuracy. However, the detailed evolutionary history and impact of tRNA modifying enzymes has not been analyzed. Using ancestral reconstruction of five tRNA modifications across 1093 bacteria, we show that most modifications were ancestral to eubacteria, but were repeatedly lost in many lineages. Most modification losses coincided with evolutionary shifts in non-target tRNAs, often driven by increased bias in genomic GC and associated codon use, or by genome reduction. In turn, the loss of tRNA modifications stabilized otherwise highly dynamic tRNA gene repertoires. Our work thus traces the complex history of bacterial tRNA modifications, providing the first clear evidence for their role in the evolution of bacterial translation.

## INTRODUCTION

Compared to the total complement of 61 sense codons, most bacterial genomes contain only 25 – 46 unique tRNA species (Grosjean, de Crécy-Lagard, & Marck, 2010). This apparent shortfall in decoding ability is mitigated by G::U wobble base pairing that allows a single tRNA to decode non–complementary codons (Crick, 1966). Additionally, specific modifications to the first anticodon base of the tRNA also contribute to wobble base pairing, sometimes reducing translational errors (Björk & Hagervall, 2014; Brierley, Meredith, Bloys, & Hagervall, 1997; Manickam, Joshi, Bhatt, & Farabaugh, 2015; Marck & Grosjean, 2002; Nasvall, Chen, & Bjork, 2004; Näsvall, Chen, & Björk, 2007; Yokoyama et al., 1985). Base modifications are carried out by several tRNA modifying enzyme (ME) pathways, modifying the wobble position of a large fraction (42%) of all tRNA species (Boccaletto et al., 2018; Grosjean et al., 2010; Machnicka, Olchowik, Grosjean, & Bujnicki, 2015). Given their potential impact on the effective tRNA pool and on translation dynamics, MEs should generally face strong selection, especially in species with a restricted tRNA repertoire. Indeed, in several bacteria, deleting modifying enzymes is lethal or otherwise deleterious for growth and translation (Björk & Hagervall, 2014; Gao et al., 2016; Shippy, Eakley, Lauhon, Bochsler, & Fadl, 2013; Wolf, Gerber, & Keller, 2002). However, in some cases MEs do not appear to be essential (Noguchi, Nishimura, Hirota, & Nishimura, 1982), suggesting strong yet variable evolutionary impact of tRNA modifications. Recently, Novoa and colleagues showed that MEs unique to the three kingdoms of life have shaped kingdom-specific tRNA pools (Novoa, Pavon-Eternod, Pan, & Ribas De Pouplana, 2012). However, the detailed evolutionary history of diverse bacterial tRNA modifications and their impact on key components of bacterial translation remain unclear.

To address these gaps, we traced the evolutionary history of the five known bacterial tRNA modifications at the first wobble base of the anticodon: (c)mnm5(s2)U (5-carboxymethylaminomethyl 2-thiouridine and its variants), cmo5U (uridine-5-oxyacetic acid), k2C (2-lysyl-cytidine), Q (Queuosine) and I (Inosine) (Björk & Hagervall, 2014). We first determined the occurrence of each modification pathway in 1093 sequenced bacteria that represent the diversity of eubacteria, and then used ancestral reconstruction to infer the major evolutionary gain or loss events in the bacterial phylogeny. To determine the causes and impacts of change in tRNA modification, we analyzed two other key components of translation: tRNA gene pools and codon bias, thought to coevolve under translational selection (Ikemura, 1985; Kanaya, Yamada, Kudo, & Ikemura, 1999). Finally, since both of these factors are in turn correlated with genomic GC content (Chen, Lee, Hottes, Shapiro, & McAdams, 2004; Hershberg & Petrov, 2009; Wald & Margalit, 2014), we also mapped genomic GC content across the bacterial phylogeny. Thus, we aimed to generate a comprehensive model of the joint evolutionary history of important components of bacterial translation – tRNA genes and modifying enzymes – and their relationship with genomic GC content.

Each modified tRNA can recognize between 2 to 4 codons. Although there are no direct measurements of the effect of tRNA modification on translation rate, the presence of a multifunctional tRNA should amplify the effective tRNA pool, reducing ribosomal waiting times for the correct tRNA and contributing to an increase in global translation speed. Modification also prevents tRNA carrying amino acids coded by two synonymous codons from recognizing other non-synonymous codons, potentially reducing mistranslation (Björk & Hagervall, 2014; Grosjean et al., 2010). Thus, tRNA modifications may affect both translational speed and accuracy. Specifically, we hypothesized that the presence of a tRNA modification should selectively favor the tRNAs that it modifies (henceforth “target” RNA). Concomitantly, selection on tRNA that are not modified by the ME (henceforth “non-target” tRNA; encoding the same amino acid) should be weakened, since their function would be redundant with the modified target tRNA. As such, we predicted that the evolutionary loss of non-target tRNA should be permissible only in the presence of a modification that allows target tRNA to decode multiple sense codons. In contrast, the evolutionary loss of a modification should be associated with a generally irreversible expansion of the tRNA repertoire to include both target and non-target tRNA. Finally, we expected that these changes in tRNA gene content should be correlated with changes in codon use as well as genomic GC content.

Our analyses indicate that the eubacterial ancestor had a large tRNA repertoire as well as most of the known bacterial tRNA modifications. However, some types of wobble were lost in multiple bacterial lineages. As predicted, ME loss was strongly associated with an expansion of the tRNA repertoire; either in conjunction with large shifts in genomic GC and associated changes in codon use, or in a GC-independent manner. We suggest that the expanded tRNA set weakened selection for MEs, allowing their repeated loss via drift. Conversely, in rare cases, a shrinking of tRNA diversity may have favored the evolution of a novel modification; alternatively, the chance evolution of a new modification may have relaxed selection on the tRNA pool. Our work represents the first systematic analysis of the joint evolutionary history of tRNA modifications, genome GC content and tRNA genes, and elucidates the role of tRNA modifications in the evolution of the bacterial translation machinery.

## RESULTS

### The evolutionary history of bacterial tRNA modifications is highly dynamic

We determined the occurrence of tRNA modifications across 1093 bacteria using homology searches for complete protein sequences of previously described MEs (see Methods). We assume that detected homologs are functional, and have the same modification function as that of the previously described enzymes. Within each modification pathway, we focused on the following enzymes that directly modify the tRNA and complete the modification (see Methods), and whose absence should effectively prevent the modification. i) MnmE-MnmG for the (c)mnm5(s2)U modification ii) CmoA-CmoB for the cmo5U modification iii) Tgt-QueA-QueG for the Q modification iv) TadA for the I modification and v) TilS for the k2C modification. Note that in our analysis, we did not consider G::U wobble pairing, which is not mediated by specific modification enzymes. We observed that except for TilS, most MEs are not found in all bacteria (Fig. 1A), suggesting that different modification pathways were gained or lost in specific lineages. The essential genes RpoB and RplA (positive controls) were respectively present in 99.5% and 99.7% of analyzed bacteria, indicating that our homology detection method resulted in very few false negatives. When multiple enzymes are necessary for a modification, they tend to co-occur in bacterial lineages (Fig. 1A). For instance, both MnmE and MnmG are absent only in Actinobacteria. Similarly, both CmoA and CmoB are primarily observed in γ-proteobacteria and ∊-proteobacteria. However, this contradicts previous reports showing that the cmo5U modification is also active in *Bacillus subtilis* and *Mycobacterium bovis* str. BCG (Chionh et al., 2016; Murao, Hasegawa, & Ishikura, 1976; Yamada, Matsugi, Ishikura, & Murao, 2005). Despite an extensive search, we could not reliably identify the reported enzymes in the sequences of any other organisms (see Methods for details). Hence, here we focus on the proteobacterial cmo5U modification. In contrast to the co-occurrence of CmoA and CmoB, for the Queuosine modification, we found that QueG is often missing even when Tgt and QueA are present. This relative evolutionary instability of QueG corroborates a recent report suggesting that Tgt alone may be enough to generate the Q modification (Zallot, Yuan, & de Crécy-Lagard, 2017).

**Figure 1:**
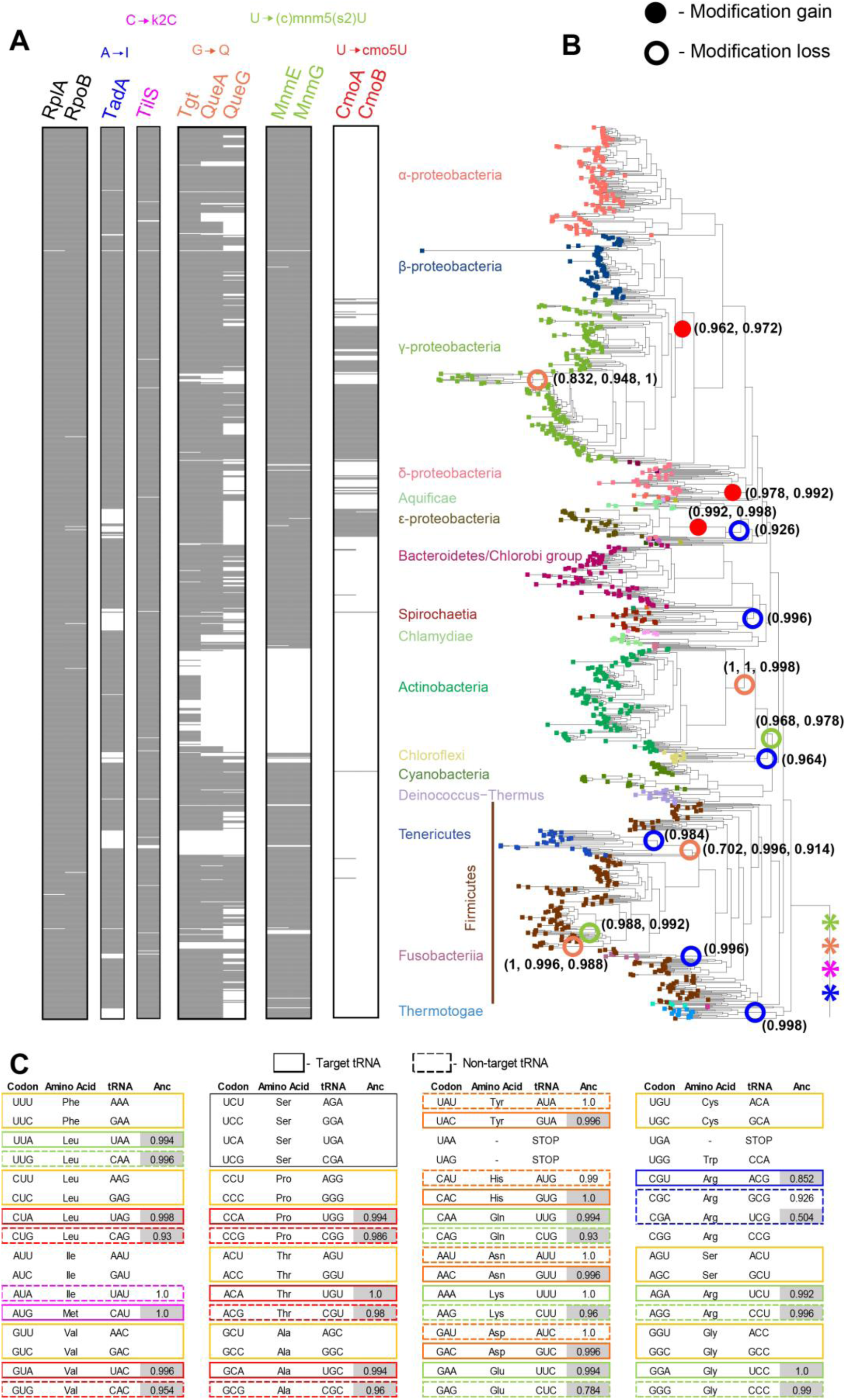
The evolutionary history of bacterial tRNA modifications. (A) Columns indicate the presence (gray) or absence (white) of various tRNA modifying enzymes as noted above each column. The leftmost column shows two essential housekeeping proteins (RpoB-RNA polymerase beta subunit and RplA-50S Ribosomal protein L1) that serve as positive controls for homology detection. (B) Bacterial phylogenetic tree (Segata et al., 2013) pruned to show the evolutionary relationships between the 1093 bacterial species used in our study. Branch tips are colored by taxonomic group. Filled and open circles respectively mark the major gain and loss of tRNA modifications (with at least 5 descendent species in the relevant lineage), as inferred by ancestral reconstruction (see Methods). Circles are colored according to the modification, as indicated in panel A. Values in parentheses indicate the posterior probability of each gain or loss event for each ME involved in the modification pathway. All other minor gain and loss events (with < 5 descendent species in the pruned phylogeny) are shown in Fig. S1. Asterisks at the root of the tree indicate the presence of the respective modification. (C) Genetic code table showing the target (solid boxes) and non-target tRNA (dashed boxes) for each modification, colored as in panel A. tRNA with a yellow border show G::U wobble base pairing and were not considered in subsequent ^16^ analyses. Serine tRNA are highlighted with a black box to indicate the ability of 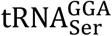 to decode all Serine codons. The column titled “Anc” indicates the presence (grey cells) or absence (white cells) of each tRNA in the eubacterial ancestor, inferred from ancestral reconstruction (see Methods). Value in each cell indicates the posterior probability of the inferred state for each tRNA.

To determine the key evolutionary changes in major tRNA modification pathways across bacteria, we performed ancestral reconstruction with stochastic character mapping for each ME (Fig. 1A; Table S1). We used a posterior probability value of ≥ 0.7 to assign the ME state at each internal node in the bacterial phylogeny. Four of the five modification systems – MnmE-MnmG, Tgt-QueA-QueG, TadA and TilS – were predicted to already occur at the root of the phylogenetic tree, i.e. were present in the eubacterial ancestor (Fig. 1B). In contrast, CmoA-CmoB were gained/evolved much later: once at the root of γ-proteobacteria, once in ∊-proteobacteria, and once in some species of δ-proteobacteria (Fig. 1B). Thus, except for the cmo5U modification, most bacterial tRNA modifications are ancient. These ancient modifications were then independently lost multiple times, with a total of 12 major losses (on branches with at least 5 descendant bacterial species in our phylogeny; Fig. 1B) and 19 minor loss events (in lineages with fewer than five descendants; Fig. S1). The only exception to this pattern of repeated ME gains/losses is the k2C modification (TilS), which appears to be evolutionarily stable across bacteria. Overall, our analysis demonstrates a highly dynamic evolutionary history of tRNA modifications that is dominated by repeated secondary losses and only a few gains (a total of 44 loss events and 13 gains across the phylogeny; Fig. S1).

### tRNA modification loss is associated with more non-target tRNAs

The early evolution of nearly all tRNA modifications suggests that the eubacterial ancestor should have been able to decode most sense codons using only a subset of all possible tRNAs. Indeed, ancestral reconstruction of tRNA genes suggests that the eubacterial ancestor did have all target ^14^ tRNAs 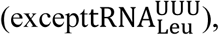 but did not possess non-target tRNAs for the Q, k2C and I modifications (Fig. 1C; note that 1C; note that the state of 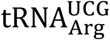 is ambiguous). Interestingly, for the (c)mnm5(s2)U modification, both target and non-target tRNA were present in the eubacterial ancestor. Thus, a subsequent loss of any of these modifications should be strongly associated with the gain (or retention, in case of (c)mnm5(s2)U) of the respective non-target tRNA. Both predictions were supported when we observed that the proportion of species that have non-target tRNA is significantly different between bacteria with vs. without a specific modification (p < 0.05, 2-sample proportions test; Fig. S2). Overall, these patterns suggest that the loss of tRNA modifications was associated with the retention or secondary gain of non-target tRNAs, and the gain of MEs was associated with the loss of non-target tRNAs.

To account for phylogenetic relationships between analysed bacteria, we compared the tRNA gene content of sister clades with contrasting modification status; i.e. following a major modification gain or loss event (see Methods; Fig. 2). For instance, since the (c)mnm5(s2)U modification was lost at the root of Actinobacteria (MnmE/MnmG enzymes, Fig. 1B), we compared the tRNA gene content of species within the Actinobacteria clade with that of species in the sister group: the Cyanobacteria-Deinococcus-Thermus (CDT) clade. Similarly, we compared the target and non-target tRNA gene copy numbers for descendent sister clades for 14 of the 15 major modification gain and loss events highlighted in Fig. 1B (we did not analyze Thermotogae, because its sister clade would consist of all other bacteria). Of these, 4 cases involving the Q modification are not informative with regard to non-target tRNAs because these are absent in all bacteria (Fig. 2F–2I). However, we describe these comparisons below since they are useful for understanding the evolution of target tRNAs.

**Figure 2:**
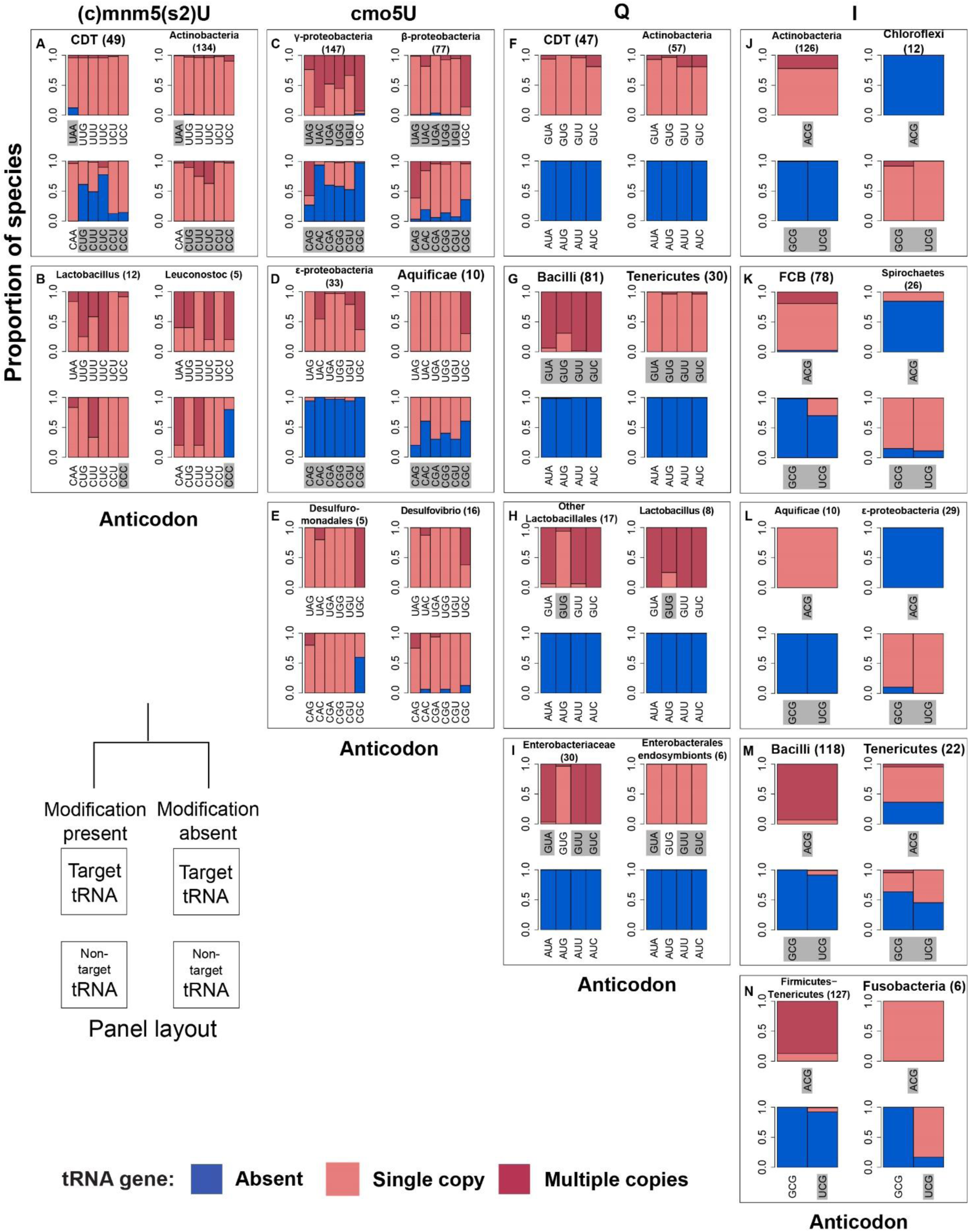
tRNA gene content associated with modification gain or loss in sister clades. Gene copy number of target and non-target tRNA within sister clades with contrasting modification status (marked in Fig. 1B). Stacked bar plots show the proportion of species within each sister clade that lack a specific tRNA (blue bars), have a single copy of the gene (light red bars), or have multiple copies (dark red bars). The total number of species analyzed in each clade are given in parentheses above the bar plots. Each panel shows data for a specific sister clade comparison; panels are grouped by modification pathway. Gray shading represents anticodons (i.e. tRNA) for which the proportion of species without the tRNA is significantly different across groups, with respect to modification presence vs. absence (p < 0.05; 2-sample test for equality of proportions with continuity correction). CDT – Cyanobacteria-Deinococcus-Thermus, FCB – Fibrobacteres-Chlorobi-Bacteroidetes.

In most of the informative sister clade comparisons (8 out of 10 cases), species were more likely to have non-target tRNAs if the modification was absent (compare bottom left vs. bottom right bar plots within each panel of Fig. 2), supporting our hypothesis that modification loss (gain) is associated with the presence (absence) of non-target tRNAs. For instance, in three out of the five comparisons made for the (c)mnm5(s2)U and cmo5U modifications, at least 5 of the 6 non-target tRNAs were present in a significantly higher proportion of species without the modification (Fig. 2A, 2C and 2D; p < 0.05, 2-sample proportions test). The two exceptions to this pattern occur in clades with relatively low sample sizes (Fig. 2B and 2E), and may potentially reflect low statistical power. Alternatively, these may be biologically interesting exceptions that require further examination. Additionally, in all five comparisons made for the I modification, at least one non-target tRNA (in most cases both) was present in significantly more species without the modification, compared to species with the modification (Fig. 2J – 2N; p < 0.05, 2-sample proportions test).

In contrast to the wide variation in the occurrence of non-target tRNAs, we found that target tRNA genes were largely stable irrespective of species’ modification status (top row in each panel of Fig. 2). Contrary to expectation, the gain or loss of the (c)mnm5(s2)U and cmo5U modifications does not appear to have any impact on target tRNAs (Fig. 2A–2E; p > 0.05, 2-sample proportions test); likely because the corresponding codons are still used in these species and cannot be decoded by non-target tRNAs. In the case of the Q modification, target tRNAs were always present; but in 2 of 4 comparisons their copy number was lower in clades that had lost the ME (Fig. 2F–2I; p < 0.05, 2-sample proportions test). Inosine presents the only evidence supporting weakened selection on target tRNAs upon ME loss. In 3 of 5 comparisons, target tRNAs were absent in nearly all species without the modification (Fig. 2J–2L). In the remaining 2 comparisons, target tRNAs were present in fewer species without modifications (Fig. 2M and 2N; p < 0.05, 2-sample proportions test). Together, these results suggest that except for the I modification, the evolution of modifications had no impact on the evolution of target tRNA genes.

### Modification status is associated with tRNA diversity, GC content and genome size

To identify the cause and impact of evolutionary changes in tRNA modifications, we compared key translation-associated genomic features of sister clades with contrasting modification status. As shown above, we found that in 11 of 14 comparisons, the tRNA diversity of sister clades was significantly different (p < 0.05, Wilcoxon rank sum test; Fig. 3 Fig. S3), indicating a strong association between the evolution of modifications and the tRNA gene pool. However, in four of these cases the direction of association opposed our initial expectation, and clades without the modification had lower tRNA diversity (Figs. 3G, 3I, 3M and 3N). In three of these cases involving Tenericutes and the endosymbiotic species from Enterobacterales (Figs. 3G, 3I and 3M), we also observed a significant reduction in the total number of tRNA genes (Fig. S3G, S3I, S3M), suggesting wide-ranging gene loss that is potentially a result of genome reduction in these clades. Indeed, previous reports have shown that reductive genome evolution in Mollicutes has led to the loss of several tRNA modifying enzymes (Grosjean et al., 2014; Yokobori, Kitamura, Grosjean, & Bessho, 2013). Thus, overall half the sister clade comparisons supported our initial prediction that the lack of modifications should be associated with greater tRNA diversity. Next, we tested whether changes in modification status and tRNA genes were also associated with changes in GC content and codon use.

**Figure 3:**
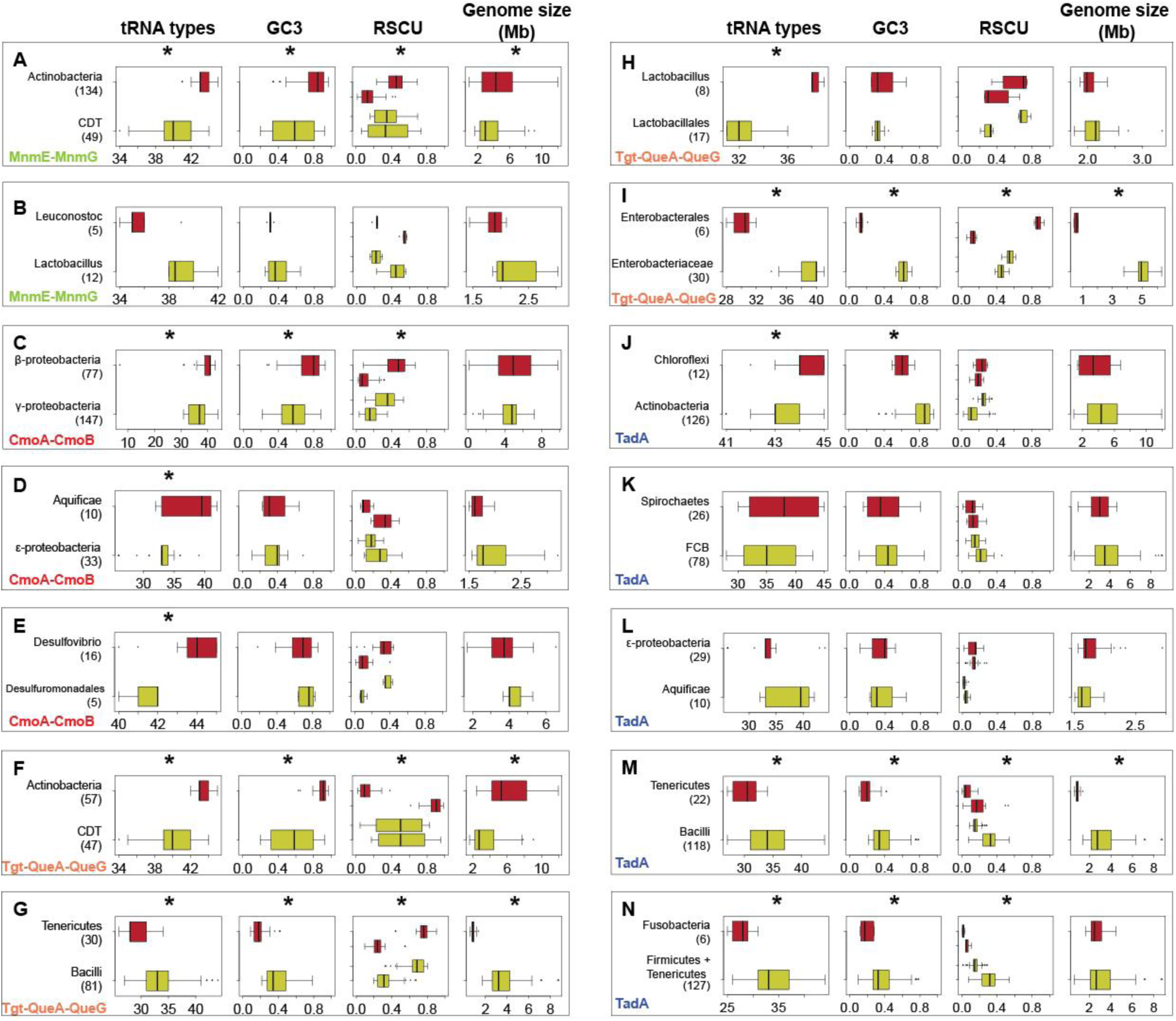
Differences in key genomic features across sister clades with contrasting modification status. Box plots show the range of values of three genomic traits, across all species within a sister clade (see Fig. 3; red boxes = modification absent; green boxes = modification present). For the Relative Synonymous Codon Usage (RSCU) panel, asterisks indicate that the difference between the distributions represented by green boxplots is significantly different than the difference between the distributions represented by the red boxplots (i.e., codon usage is more biased in one case; p < 0.05; Wilcoxon rank sum test with p-values correction for multiple comparisons). For tRNA types, GC3 (GC content of the third codon base) and genome size, asterisks indicate a significant difference in distributions across the sister clades (p < 0.05; Wilcoxon rank sum test with p-values correction for multiple comparisons).

In most cases showing altered tRNA diversity (8 out of 11), we found that the change in diversity was also associated with significant shifts in GC content, so that species in clades without the tRNA modification typically had more extreme GC3 values (i.e. GC content in the third base of codons, Fig. 3; p < 0.05, Wilcoxon rank sum test; we observed similar results with genomic GC content, Fig. S3). In seven of these eight cases, species without the modification also had significantly more biased codon use (measured as relative synonymous codon use RSCU for all proteins; p < 0.05, Wilcoxon rank sum test; we observed similar results for RSCU of highly expressed ribosomal proteins; Fig. S3). Finally, in five of the eight cases we observed significant differences in genome size across sister clades (Fig. 3). Of these, in the two comparisons that involved Actinobacteria (Figs. 3A and 3F) we observed that larger genomes were associated with the absence of the modification. As mentioned above, in the other three cases involving Tenericutes and Enterobacterales (Figs. 3G, 3I and 3M), the lack of modification may reflect overall genome reduction. Note that in every comparison where codon use and genome size were different, GC content was also skewed significantly. Thus, to test the overall effect of genome size and GC content on modification status, we carried out a phylogenetic regression. We found that both GC content and genome size had a significant effect on the presence/absence of the Q modification alone (Fig S4). These results suggest that except for the Q modification, modifications were lost in only a few instances of shifts in GC content. Thus, GC content may be the major ultimate driver of changes in tRNA gene content and the subsequent loss of modification may generate selection to fix these changes. Altogether, three of the 14 major changes in modification status were not associated with significant shifts in either tRNA or GC content, and remain unexplained. These included one instance of the loss of the (c)mnm5(s2)U modification (Fig. 3B) and two losses of the I modification (Figs. 3K–L).

Finally, to quantify the evolutionary impact of tRNA modifications on tRNA gene pools, we simultaneously mapped changes in tRNA genes copies, MEs, and GC content on the bacterial phylogeny. As expected, following major GC shifts, non-target tRNA showed several gain/loss events whereas target tRNA were more stable (Figs. 4–6). As suggested by our sister clade comparison, in some cases modification gain or loss was also preceded by a GC shift (8 out of 14 independent lineages). Moreover, a phylogenetic regression to test the impact of GC and MEs on tRNA genes revealed that GC content had the strongest impact on the dynamics of non-target tRNA genes (Fig. 7). However, as observed in the sister clade comparison, the phylogenetic regression and mapping also reveal clear instances of GC-independent impacts of modification status on tRNA genes (Figs. 4–7). The strongest impact of tRNA modification is observed the case of the Inosine, where the target tRNA was present in all species when the modification was present (Fig. 6); and the non-target tRNA were absent in most of these species (Fig. 2D). On the other hand, the loss of target tRNAs and presence of non-target tRNAs is strongly associated with the loss of the Inosine modification (Fig. 2D, Fig. 6, Fig. 7). Thus, the bacterial tRNA repertoire is strongly affected by evolutionary changes in GC content as well as tRNA modifying enzymes.

**Figure 4:**
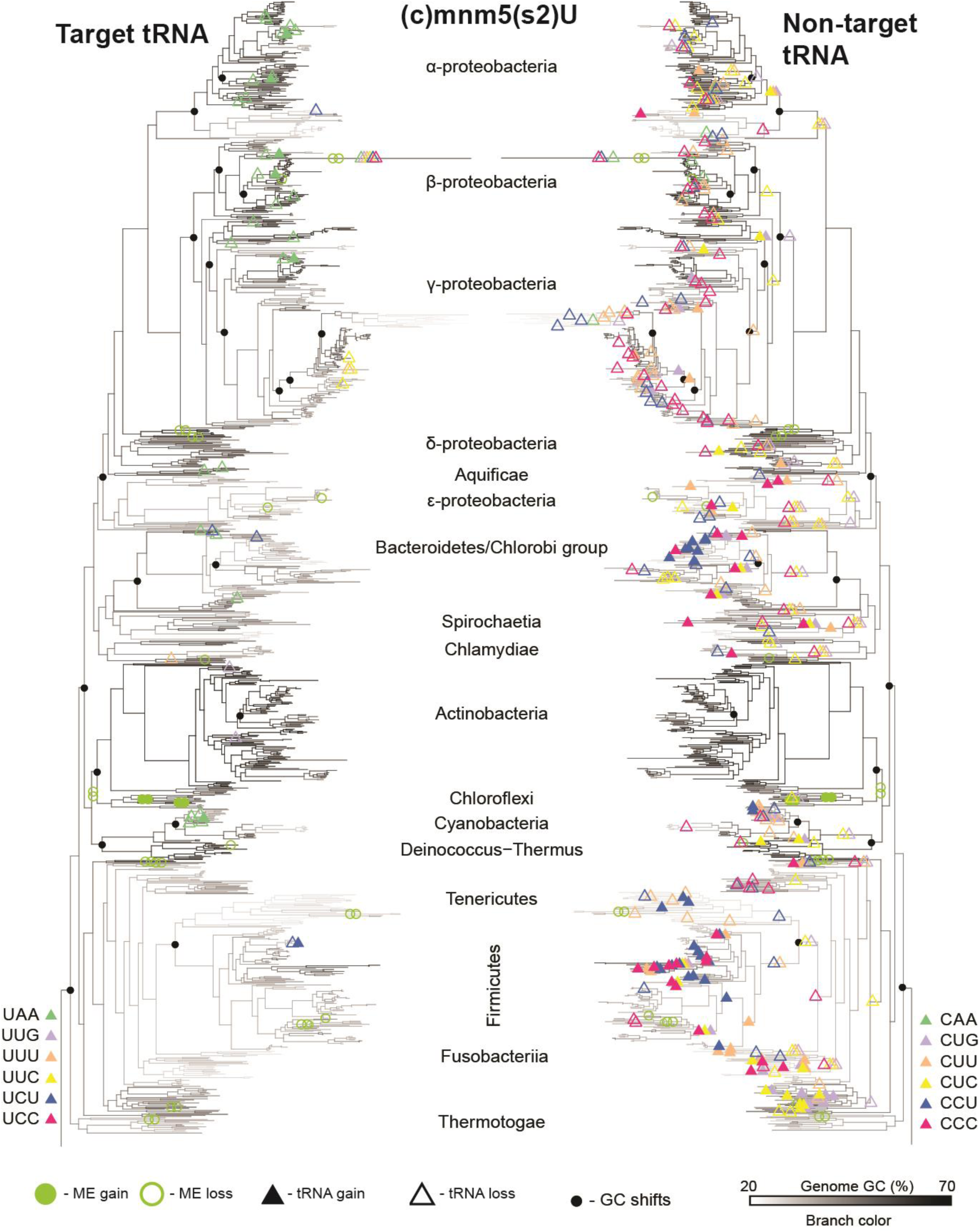
Evolutionary dynamics of tRNA genes, MEs and GC content for the (c)mnm5(s2)U modification. Branches are colored according to the inferred GC content for each branch, with white indicating low GC% and black indicating high GC%. Nodes that showed the largest GC shifts are indicated with black circles. Colored open and closed circles respectively indicate the loss and gain of each ME involved in the modification. Open and closed triangles indicate the loss and gain of tRNA genes relevant to the (c)mnm5(s2)U modification, with target tRNA indicated on the tree on the left and non-target tRNA indicated on the mirror tree on the right.

**Figure 5:**
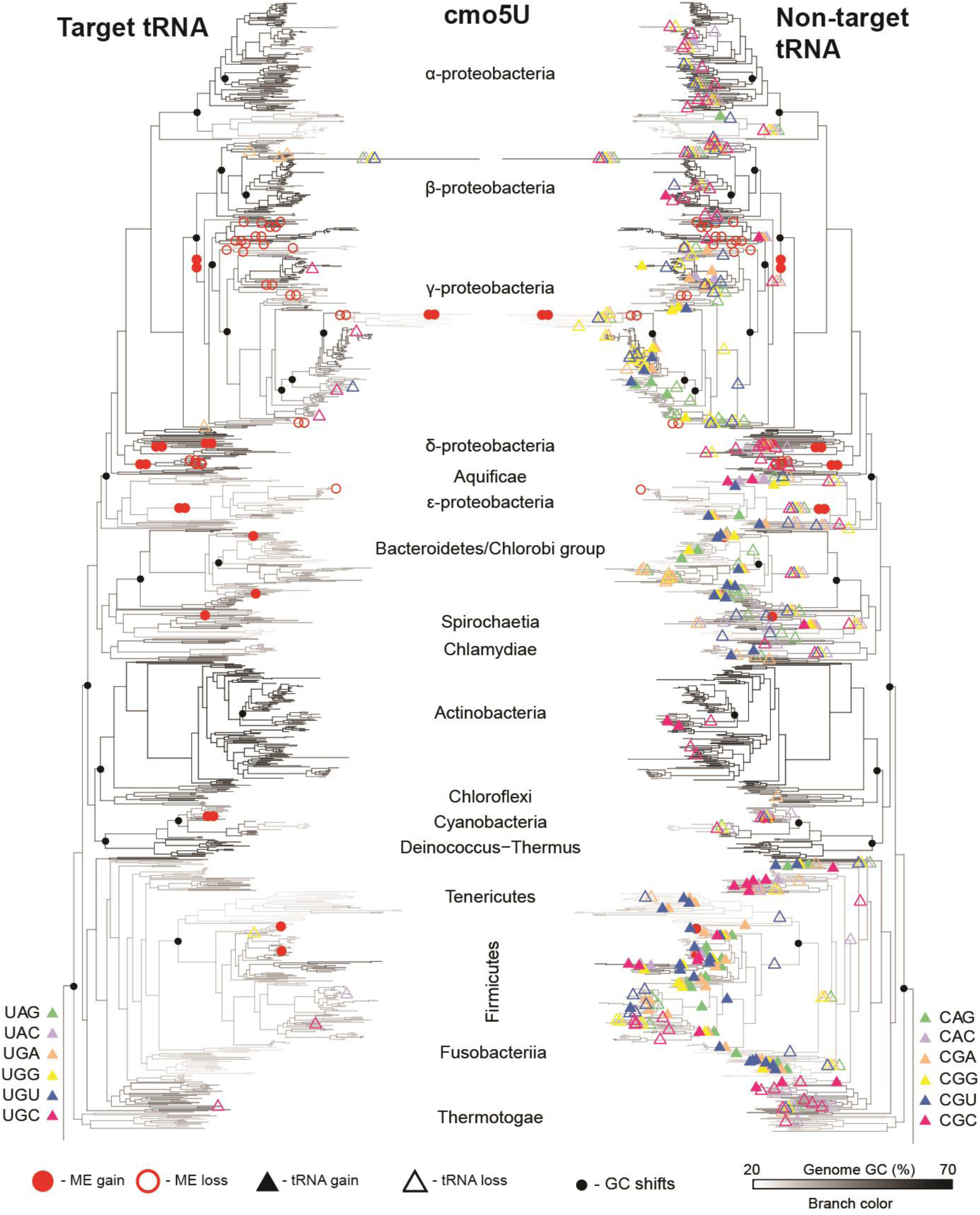
Evolutionary dynamics of tRNA genes, MEs and GC content for the cmo5U modification. Branches are colored according to the inferred GC content for each branch, with white indicating low GC% and black indicating high GC%. Nodes that showed the largest GC shifts are indicated with black circles. Colored open and closed circles respectively indicate the loss and gain of each ME involved in the modification. Open and closed triangles indicate the loss and gain of tRNA genes relevant to the cmo5U modification, with target tRNA indicated on the tree on the left and non-target tRNA indicated on the mirror tree on the right.

**Figure 6:**
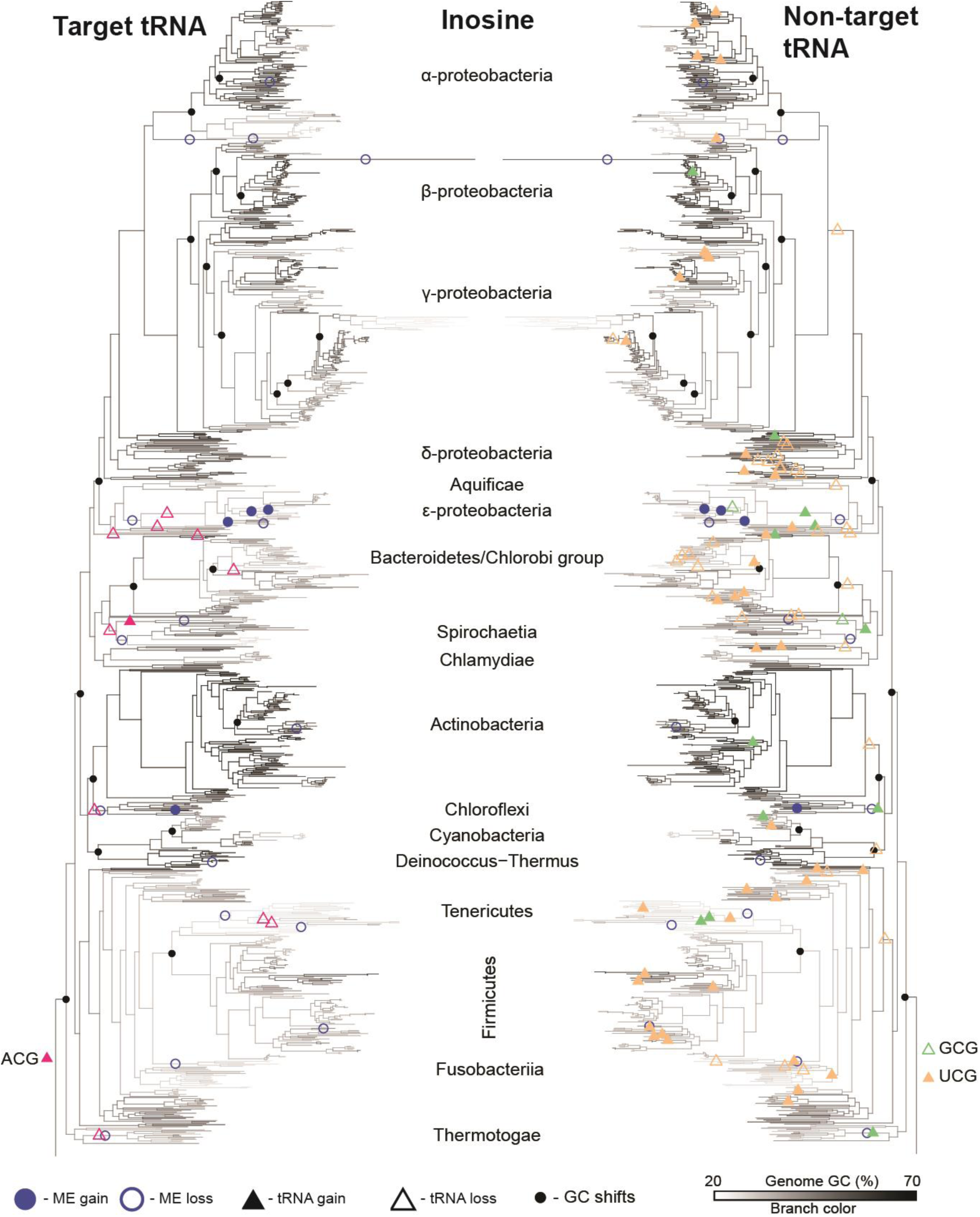
Evolutionary dynamics of tRNA genes, MEs and GC content for the I modification. Branches are colored according to the inferred GC content for each branch, with white indicating low GC% and black indicating high GC%. Nodes that showed the largest GC shifts are indicated with black circles. Colored open and closed circles respectively indicate the loss and gain of each ME involved in the modification. Open and closed triangles indicate the loss and gain of tRNA genes relevant to the I modification, with target tRNA indicated on the tree on the left and non target tRNA indicated on the mirror tree on the right.

**Figure 7:**
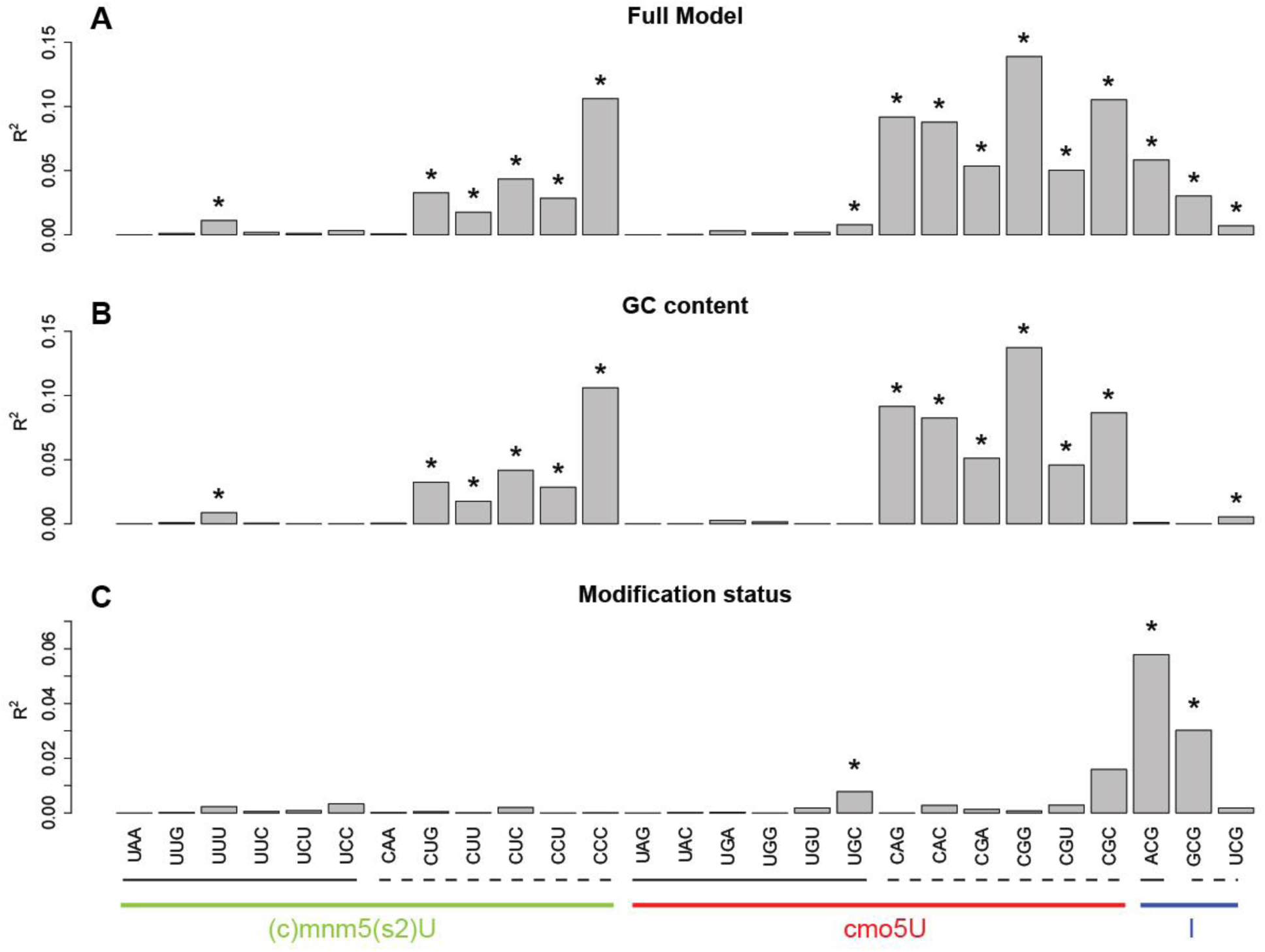
Results of phylogenetic regressions for factors affecting tRNA gene presence or absence. Bar plots show the R^2^ value for the phylogenetic regression for presence/absence of target and non-target tRNAs for the (c)mnm5(s2)U, cmo5U and I modifications. Results are shown for three different models: (A) Full model, tRNA presence/absence ~ Modification status (i.e. presence/absence) * GC content. (B) tRNA presence/absence ~ GC content (C) tRNA presence/absence ~ Modification status (i.e. presence/absence). Asterisks indicate a significant impact of the focal trait on the presence of the tRNA gene (see Methods). Target tRNAs are highlighted with a solid underline, and non-target tRNAs with a dashed underline. Note the different y-axis scale in panel C.

## DISCUSSION

Over 50 years ago, Francis Crick described the phenomenon of wobble base pairing where one tRNA molecule can recognize multiple codons (Crick, 1966). Since then, a large body of work has developed a detailed understanding of the enzymatic pathways that generate extended wobble pairing via base modifications of tRNA anticodons (reviewed in (Björk & Hagervall, 2014)). Given their critical role in translation, modifying enzymes are expected to evolve under strong selection, and in turn influence the evolution of other components of the translation machinery (Grosjean et al., 2010). Here, we trace the evolutionary history of tRNA modifications across the bacterial phylogeny, showing that the eubacterial ancestor already had nearly all known modification pathways. Several major bacterial clades subsequently lost the modifications, often accompanied by increasing tRNA diversity and shifts in genome GC content and codon use. These results are consistent with the hypothesis that an expanded tRNA pool gradually weakens selection on MEs, allowing their loss through drift. However, once the modification is lost, bacteria likely face strong selection to retain the expanded tRNA pool. In contrast, if GC shifts shrink the tRNA gene pool, selection would strongly favor the innovation or retention of tRNA modifications. We speculate that in Proteobacteria, such a process may have led to the evolution of the novel cmo5U modification. Alternatively, Proteobacteria may have evolved the new modification by chance, secondarily weakening selection on the non-target tRNAs. Interestingly, we also uncovered instances where changes in tRNA or GC content could not explain the evolution of tRNA modifying enzymes. In some of these cases, we suspect that overall genome reduction may have led to the loss of the modification; but three major cases of modification loss remain unexplained (loss of (c)mnm5(s2)U in Leuconostoc and I in Spirochaetes and ∊-proteobacteria). Nonetheless, our results strongly support the conclusion that ME evolution was dominated by frequent losses driven by weakened selection. Note that reversing the direction of causality in our arguments would not make logical sense: the loss of non-target tRNAs cannot generate selection favoring the loss of tRNA modifications, and there is no reason to expect that the gain of a modification enzyme should selectively favor the retention on non-target tRNAs. Similarly, it is difficult to imagine how evolutionary gains or losses of tRNA modifications could precipitate genome-scale changes in GC content or genome size. Thus, our systematic phylogenetic analysis clarifies the contrasting roles of strong vs. weak selection acting on tRNA modifications, and on the evolutionary history of key components of bacterial translation.

Our results were generally consistent with the prediction that a lack of specific tRNA modifications should result in strong positive selection favoring the relevant non-target tRNAs. However, we also observed some instructive exceptions to this pattern. For instance, comparing across sister clades, we found that ~33% of the species that do not have the cmo5U modification also lack at least one non-target tRNA. This should pose a problem because codons recognized by the missing non-target tRNAs cannot be decoded efficiently. However, we observed that these codons were significantly depleted in these genomes, compared to species that retained the non-target tRNA (Fig. S5). Thus, these relatively rare codons may not pose a major problem even if they are inefficiently translated. It is also possible that these codons are decoded by the respective U-starting target tRNAs that can pair with A, G or U ending codons (Grosjean et al., 2010). This scenario is especially likely in species where the Q modification is absent. Recall that all bacteria lack the non-target tRNA for this modification; hence species without the Q modification should be unable to translate many codons. We hypothesize that in species where both the Q modification as well as the non-target tRNA are missing, there is an as yet undiscovered G::U wobble base pairing between the target tRNA and codons complementary to the non-target tRNA in these species.

Our analysis quantifies the role of specific tRNAs in the evolutionary dynamics of bacterial translation, both as drivers of selection on tRNA modifications and as a means to rapidly respond to shifts in genomic GC content. Specifically, we found that target tRNAs are evolutionarily largely stable (except for those modified by Inosine), whereas non-target tRNAs are often gained or lost in lineages following a GC shift. The strong phylogenetic correlation between non-target tRNAs and genomic GC content echoes the previously reported association between GC content and “auxiliary” tRNAs (tRNAs with a high number of gains and losses) defined by Wald and Margalit (Wald & Margalit, 2014). The nearly ubiquitous presence of target tRNAs may reflect strong positive selection, since the function of target tRNA cannot be carried out by any other tRNA via wobble base pairing rules. Thus, target tRNA may be essential even if their complementary codons are used relatively rarely.

In summary, we propose that the evolution of diverse bacterial tRNA modification enzymes has been strongly influenced by changes in tRNA gene pools and genomic GC content, as well as by genome reduction in specific lineages. Our results support the hypothesis that these changes weakened selection on MEs and allowed their loss through drift. Conversely, we hypothesize that the loss of non-target tRNA may have generated strong selection for the innovation of the cmo5U modification in γ-proteobacteria and ε-proteobacteria. It is also possible that the tRNA loss succeeded only in the wake of the chance innovation of the cmo5U modification. These patterns generate a number of broad, testable predictions about the future evolutionary trajectory of translational components in various bacterial lineages. For instance, MEs may be lost in GC-rich bacteria without major consequences for translation; such losses should evolutionarily fix the expanded tRNA repertoire; and MEs should face strong positive selection in AT-rich clades with low tRNA diversity. Specifically, in AT rich bacteria, the combination of MEs and largely target tRNAs should confer higher fitness than a full tRNA set without MEs. Our analysis also shows that multiple key components of bacterial translation have evolved in close association with genomic GC content. Although the evolutionary and ecological forces driving GC shifts and GC-independent changes in tRNA diversity remain unclear, our analyses highlight the rich evolutionary dynamics of wobble pairing, and more broadly, bacterial translation.

## METHODS

### Phylogenetic tree, genomes and tRNA gene data

All analyses were carried out in R (R Development Core Team, 2015) using the phytools (Revell, 2012) and seqinr (Charif, Thioulouse, Lobry, & Perrière, 2005) packages, unless otherwise specified. The R scripts for all analyses are available at https://github.com/gauravdiwan89/me_evolution_project. We used the bacterial phylogeny generated by Segata and co-workers (Segata, Börnigen, Morgan, & Huttenhower, 2013). This phylogeny was generated using an alignment of the 400 most conserved protein sequences across all prokaryotes. We downloaded the phylogeny (https://bitbucket.org/nsegata/phylophlan/wiki/bs_tree.reroot.nwk) and removed all archaea, as well as bacteria whose genomes have not been fully sequenced. We pruned the tree further using a distance based pruning function. Briefly, when closely related species had a pairwise phylogenetic distance ≤0.03, we randomly sampled only one of these species. This allowed us to remove the excessive representation of some highly sequenced genomes, such as several *Escherichia coli* strains. This led to the final set of 1093 bacterial genomes that we used in our subsequent analyses.

We downloaded the tRNA gene copy numbers for each of these genomes from the Genomic tRNA database (Chan & Lowe, 2009). For genomes where this information was not available, we determined the tRNA gene copy numbers by running the program tRNAscan-SE (Lowe & Eddy, 1996) using the default parameters for bacterial genomes. The information for each of these genomes was represented using different identifiers in the various databases that we used. This information is consolidated in Table S1, consisting of: i) species name according to NCBI taxonomy, ii) NCBI accession number, iii) Phylogenetic tree tip label in the tree, iv) IMG database identifier and v) Genomic tRNA database abbreviation.

### Choice of crucial tRNA MEs and non-target tRNA

The (c)mnm5(s2)U and Q modifications are created by elaborate ME pathways. For instance, our definition of the (c)mnm5(s2)U modification includes three different modifications: mnm5U, cmnm5U and cmnm5s2U, each of them acting on different isoaccepting tRNA. These modifications have unique MEs that are involved in the last steps of the pathway. MnmE and MnmG are the common enzymes for each of these modifications and they carry out the first step of the modifying the Uridine base (Björk & Hagervall, 2014) and references therein. Thus, we considered MnmE-MnmG to be crucial, and other enzymes such as MnmC, MnmA, MnmH and trmL as auxiliary. The Q base is first biochemically synthesized as Q_1_ in the bacterial cell using the enzymes FolE, QueC, QueD, QueE and QueF. Subsequently, the Tgt enzyme replaces Guanosine in the anticodon stem loop with Q_1_ and QueA and QueG convert it to Q ((Björk & Hagervall, 2014) and references within). A recent report has suggested the possibility of Tgt alone carrying out the Q modification in the absence of QueA and QueG, assuming the existence of a Q salvage pathway (Zallot et al., 2017). However, there is no evidence for this phenomenon across most bacteria from our analysis; hence, we considered Tgt, QueA and QueG as the crucial enzymes for the Q modification.

The cmo5U modification exists in tRNA molecules for four box codons (Leucine, Valine, Serine, Proline, Threonine and Alanine) and therefore there were three possible non-target tRNA in each case. However, tRNA with GNN anticodons (where N is any nucleotide) in each of these codon boxes can also recognize NNU codons via G::U wobble pairing. Thus, we only focused on the non-target tRNA that have CNN anticodons (indicated in Fig. 1C).

### Homology search for tRNA MEs

We determined the presence/absence of all known tRNA MEs in the 1093 bacterial genomes using Hidden Markov Model (HMM) searches (S. Eddy, 1998). First, we downloaded all available reviewed protein sequences for each of the MEs from the UniProt-KB database. We created a multiple sequence alignment using the t-coffee program (Notredame, Higgins, & Heringa, 2000) with default parameters. We used this alignment to build an HMM profile, and then used the “hmmsearch” command in the HMMER suite (S. R. Eddy, 2009) to detect the presence/absence of each ME. We used the amino acid sequences of all proteins from the 1093 genomes (downloaded in November 2016 from the NCBI ftp site) as the sequence search space, and set all other parameters to default. Each search generated a table with accession numbers of potential homologous hits with a corresponding e-value and bit-score. There is no consensus in the field with respect to setting an e-value cut-off to detect true homologs. Here, we present a slightly informative method of deciding e-value cut-offs for detecting true homologs. To infer presence/absence with precision, we implemented a dynamic e-value cut-off to eliminate spurious hits. We did this by plotting the number of species in which the ME would be detected as we increased the e-value cut-off (Fig. S6). We observed stable plateaus where the number of species in which an ME was detected did not change despite changing the e-value cut-off, suggesting that these represent robust e-value thresholds. Thus, for each enzyme we set an enzyme-specific e-value cut-off as the first e-value at the beginning of the most stable plateau (see Fig. S6). For each ME, homologous hits that had e-values lower than this cut-off were considered true homologues.

Previous reports have experimentally shown that the cmo5U modification occurs in *Bacillus subtilis* and *Mycobacterium bovis* str. BCG (Chionh et al., 2016; Murao et al., 1976; Yamada et al., 2005). However, in our initial analysis we did not find CmoA and CmoB homologs in these or related species. A closer inspection of the *Bacillus subtilis* genome also did not reveal these enzymes, as indicated in a previous report (Grosjean et al., 2014). Instead, there were a few entries for these enzymes in the UniProt database (each with a poor annotation score). We therefore tested whether these enzymes are true homologs of the proteobacterial CmoA and CmoB, using homology detection (using ‘jackhmmer’). We observed that only the query organisms showed the presence of these enzymes (data not shown). A survey of the SEED database (Overbeek et al., 2005) also revealed the absence of CmoA and CmoB in all but one *Bacillus* species. When we carried out a blastp analysis using CmoA and CmoB enzymes from *Escherichia coli* in *Mycobacterium bovis* str. BCG, we observed no significant hits for CmoA and no hits for CmoB with e-values < 5e-3. Thus, although the modification may exist in several species, we could not reliably identify homologs of the proteobacterial CmoA and CmoB in other bacteria.

### Ancestral reconstruction of tRNA modifications, genomic GC content, and tRNA genes

We built a presence-absence matrix for each ME and used the binary state of each ME to infer gain or loss events along the phylogeny. We used the stochastic character mapping (Bollback, 2006) function in the phytools package in R, with the following parameters: transition rate matrix determination method – Bayesian MCMC, prior distribution on the root node of the tree – estimated, number of simulations – 500. We fixed the number of simulations such that the posterior probability that an enzyme is present for 100 randomly sampled nodes did not change (Fig. S7). We used the enzyme with the most variable presence/absence state (queG) for this analysis, and assumed that calculations for all other enzymes would be more stable. The mapping resulted in 500 enzyme states at each node of the tree. We determined probabilities of each state at each node and assigned the state that had a posterior probability of ≥ 0.7 (Fig. S8) as the state of the enzyme at that node. In case the probability was < 0.7, we assumed the same state at the node as the preceding node. To determine gain and loss of tRNA modification pathways, we marked the branch that succeeded the gain or loss of all MEs for a given modification as the focal branch. Only these events are described in this study; however, all data for each enzyme are shown in Fig. S1.

We determined the GC content at each ancestral node in the bacterial phylogeny using the StableTraits program (Elliot & Mooers, 2014) with 10 million iterations and all other parameters set to default. We then determined 20 nodes whose two immediate descendants had the largest difference in GC. Each of these nodes had at least 10 descendant species. These nodes representing major GC shifts are shown on the phylogeny along with the ancestral GC content (Figs. 4–6).

We determined the evolutionary history of tRNA gains and losses using a similar method as described for MEs. However, in this case we carried out the ancestral reconstruction separately for each bacterial family, which allowed us to account for family-specific rates of evolution of tRNA genes. We used the ancestral state of each tRNA at the root of each bacterial family to infer the tRNA gene content of the eubacterial ancestor. Briefly, we converted tRNA gene copy numbers for all species into a binary vector by changing the state of tRNA genes with multiple copies to “1” i.e. presence. We then carried out ancestral reconstruction using stochastic character mapping and noted evolutionary transitions on the phylogenetic tree as before.

### Identifying sister clades with contrasting modifying enzyme status

We identified clades where crucial enzymes for each tRNA modification were gained or lost, and determined the closest sister clade where the state of modification was opposite to that of the focal clade. We determined the sister clade agnostic of its taxonomic classification. We checked if there were at least 5 descendant extant species in this clade for each gain or loss event. If this criterion was not met, we picked the next closest sister clade neighboring the common ancestor of the above two clades. Within each clade, we excluded species that had an opposite state for any of the MEs involved in each modification. We assigned the identity of the comparison group using the lowest level of taxonomic classification that encompassed the focal species.

### Phylogenetic regression

We carried out phylogenetic regression to test the impact of GC content and modifcations on tRNA presence/absence, using the Maximum Likelihood Continuous Regression program in BayesTraits, similar to the method used by Wald and Margalit (Wald & Margalit, 2014). We estimated the lambda parameter for each case and used default values for all other parameters. For each tRNA, we first tested the full model: tRNA presence/absence ~ GC content*tRNA modification status. To determine the impact of each explanatory variable, we compared the likelihood of the full regression model to the likelihood of the same regression when the R value was set to zero. For reduced models containing only GC content or modification status as the explanatory variable, we compared the likelihood of the regression of the reduced model with the likelihood of the regression of the full model. When these likelihoods were comparable (p > 0.05; Likelihood Ratio test), we inferred that the reduced model was sufficient to explain tRNA presence/absence, and hence had a significant impact on the evolution of the tRNA. We used the same method for testing the impact of GC content and genome size on modification status (Fig. S4).

## ACKNOWLEDGEMENTS

We thank Saurabh Mahajan for help with analyses; and Aparna Agarwal, Shyam Buddh, Vrinda Ravi Kumar, Saurabh Mahajan and Laasya Samhita for constructive comments on the manuscript. This work was funded by the National Centre for Biological Sciences (NCBS-TIFR) and grants from the Department of Science and Technology, India (INSPIRE Faculty award, IFA-1 LSBM-64) and the Council for Scientific and Industrial Research, India (37(1629)/14/EMR-II). GDD acknowledges a Senior Research Fellowship from the University Grants Commission, India.

